# Longitudinal cellular and humoral immune responses following Covid-19 BNT162b2-mRNA-based booster vaccination of craft and manual workers in Qatar

**DOI:** 10.1101/2024.12.10.627725

**Authors:** Remy Thomas, Ahmed Zaqout, Bakhita Meqbel, Umar Jafar, Nishant Vaikath, Abdullah Aldushain, Adviti Naik, Hibah Shaath, Neyla S Al-Nakl, Abdi Adam, Houda Y Ali Moussa, Kyung C Shin, Rowaida Z Taha, Mohammed Abukhattab, Muna A Almaslamani, Nehad M Alajez, Abdelilah Arredouani, Yongsoo Park, Sara A Abdulla, Omar MA El-Agnaf, Ali S Omrani, Julie Decock

## Abstract

**Introduction:** In March 2020, the rapid spread of SARS-CoV-2 prompted global vaccination campaigns to mitigate COVID-19 disease severity and mortality. The 2-dose BNT162b2- mRNA vaccine effectively reduced infection and mortality rates, however, waning vaccine effectiveness necessitated the introduction of a third vaccine dose or booster. To assess the magnitude and longevity of booster-induced immunity, we conducted a longitudinal study of SARS-CoV-2 specific cellular and humoral immune responses among Qatar’s vulnerable craft and manual worker community. We also investigated the impact of prior naturally acquired immunity on booster vaccination efficacy.

**Methods:** Seventy healthy participants were enrolled in the study, of whom half had prior SARS-CoV-2 infection. Blood samples were collected before and after booster vaccination to evaluate immune responses through SARS-CoV-2 specific ELISpots, IgG ELISA, neutralization assays, and flow cytometric immunophenotyping

**Results:** T cell analysis revealed increased Th1 cellular responses, marked by enhanced IFN-γ release, in recently infected participants, which was further enhanced by booster vaccination for up to 6-months. Furthermore, booster vaccination stimulated cytotoxic T cell responses in infection-naïve participants, characterized by granzyme B production. Both natural SARS-CoV-2 infection and booster vaccination induced robust and durable SARS-CoV-2 specific humoral immune responses, with high neutralizing antibody levels. Prior natural infection was also linked to an increased number of class-switched B cells prior to booster vaccination.

**Conclusion:** These findings underscore the importance of booster vaccination in enhancing anti-viral immunity across both infection-naïve and previously infected individuals, enhancing distinct arms of the anti-viral immune response and prolonging naturally acquired immunity.

## INTRODUCTION

The emergence of the novel human pathogen, severe acute respiratory syndrome coronavirus 2 (SARS-CoV-2) led to a global pandemic in 2019. SARS-CoV-2 is believed to have originated from bats in the Wuhan province in China and rapidly spread to humans through carrier animals and human-to-human transmission (1). The virus causes Coronavirus Disease 19 (COVID-19), a highly infectious disease with a range of clinical manifestations from asymptomatic infections to severe respiratory failures. Among infected individuals, 10-15% required hospitalization, with 15-20% of those needing intensive care, (2, 3). Similar to other coronaviruses, SARS-CoV-2 is a single-stranded, positive-sense RNA that encodes several structural proteins including the spike (S), envelope (E), membrane (M) and nucleocapsid (N) proteins which are essential for viral entry, replication and assembly (4). The spike protein in particular plays a critical role in the virus’ pathogenesis by binding to the angiotensin-converting enzyme 2 (ACE2) receptor on host cells, facilitating viral entry (5). Therefore, mutations in the spike protein can greatly impact the transmissibility of the virus, as observed with the SARS-CoV-2 variants of concern (VOC) Alpha, Delta and Omicron (6). As SARS-CoV-2 evolved, numerous genetic variants emerged, displaying distinct phenotypic characteristics including differences in transmission rate, disease severity and ability to escape immunosurveillance. Moreover, the duration of infectiousness evolved, with individuals infected with the Omicron variant exhibiting an earlier onset of infectiousness compared to those infected with the Delta variant, allowing for more rapid transmission (7).

To reduce the global health burden of COVID-19, public health measures and global vaccination campaigns were rapidly implemented. The World Health Organization (WHO) approved a total of 21 vaccines, including vaccines based on inactivated viruses, protein, recombinant adenovirus, DNA, and messenger RNA (mRNA) (8). Despite the major efforts in developing these COVID-19 vaccines, vaccination with a single dose of the Pfizer/BioNTech (BNT162b2) and Moderna (mRNA-1273) mRNA vaccines may not provide sufficient long-lasting protection (9–11). In contrast, a two-dose regimen significantly improved vaccine effectiveness (VE), with the Pfizer/BioNTech vaccine achieving over 90% effectiveness shortly after the second dose and Moderna over 80% (12–16). Nevertheless, a meta-analysis of 18 studies revealed waning immunity following two-dose vaccination with BNT162b2 (Pfizer-BioNTech), mRNA-1273 (Moderna) and Ad26.COV2.S (Janssen) against SARS-CoV-2 infection, as demonstrated by a decrease in pooled mean VE from 83% at one month post-vaccination to 22% at five months, with a sharp decline after 100 days following full vaccination (17, 18). Furthermore, a large study of approximately 10.6M individuals receiving a two-dose mRNA vaccine regimen reported a decline in BNT162b2 and mRNA-1273 VE going from respectively 94.5% and 95.9% at two months after the first dose to 66.6% and 80.3% at seven months (18). Notably, one study investigated the waning of the two-dose BNT162b2 and mRNA-1273 VE with bias correction for pre-vaccination prevalence of naturally acquired immunity in order to evaluate differences in susceptibility attributable to vaccination only (19). Without bias correction, they found a pooled VE of 91.3% at 14 days post second-dose which declined to 50.8% at 7 months post vaccination. Similarly, bias-corrected VE at 7 months post-vaccination reached 53.2%, suggesting that waning of protection by vaccination is not impacted by pre-vaccination naturally acquired immunity. In line with a waning vaccine effectiveness, two doses of the BNT162b2 vaccine were shown to elicit humoral and adaptive immune responses for up to five months after the first dose (20). Analysis of six healthy, adult vaccine recipients showed an early increase in anti-spike antibody responses after the first dose (day 20), followed by a second increase after the second dose (day 34) and subsequent decline 150 days after the first dose. Notably, S1-specific T cell responses mirrored this pattern, with the second dose enhancing T cell responses in all six recipients, four of whom exhibited detectable responses up to five months after the first dose. Furthermore, it has been postulated that the waning protection of the two-dose vaccination regimen may be due to a decline in immunity in addition to the emergence of the Delta variant. For example, different two-dose BNT162b2 and mRNA-1273 vaccine effectiveness values were reported in epochs during which the Alpha VOC was the most prevalent (VE mRNA-1273=93.7% and VE BNT162b2=85.7%) compared to epochs where the Delta VOC was more present (VE mRNA-1273=75.6% and VE BNT162b2=63.5%) (21).

Together, the waning of protection and emergence of VOCs prompted the need for booster doses to restore immunity and provide better protection against COVID-19 VOCs. Previous findings from Qatar showed a sharp drop in BNT162b2 VE to below 40% at 181-270 days following the second dose, whereas administration of a third or booster dose was associated with a VE of approximately 80% (22). Similarly, three-dose BNT162b2 and mRNA-1273 VE values at 4-11 months post second dose have been shown to be comparable to VE values of 2-dose regimens at 1 to 2 months following full vaccination of a predominantly white, non-Hispanic population (23). Anti-spike antibody levels in recipients of two doses of a mRNA vaccine (BNT162b2, mRNA-1273) have been reported to peak at 90 days after the first dose, followed by a decline from day 90-180 (24). Following the third dose, recipients’ antibody levels increased almost 2.5-fold compared to the levels observed at day 90, after which a gradual decline was observed from day 251 to day 535 to levels which remained higher than the peak at day 90. Overall, administering a third dose of either BNT162b2 or mRNA-1273 has been shown to enhance antibody level persistence with a slower decline compared to two-dose regimens. Notably, booster BNT162b2 vaccination resulted in higher antibody avidity

In this study, we examined the effects of the third dose of the BNT162b2 vaccine on the dynamics of cellular and humoral immune responses in craft and manual workers (CMWs) in Qatar. Previous studies highlighted that this community, constituting approximately 80% of Qatar’s population, experiences a higher rate of infection which is likely due to multiple factors including overcrowded living and working conditions and educational challenges (26–29). Given this higher vulnerability, we sought to investigate how administering a third vaccine dose may affect immune responses in this population and whether prior natural infection influences vaccine-induced immunity. Using diverse approaches, we observed that administering a third dose of the BNT162b2 vaccine effectively induced both cellular and humoral immune responses in our CMW population, while also enhancing the pre-existing immunological memory in participants with a prior natural SARS-CoV-2 infection.

## MATERIALS AND METHODS

### Study population

A total of 70 healthy adults from the CMW community in Qatar were included in the study. All participants received two doses of the BNT162b2 mRNA vaccine and were scheduled to receive the third dose at the Communicable Disease Center in Qatar between May 25, 2022 and July 4, 2022. None of the participants tested positive for SARS-CoV-2 infection within 4 weeks prior to the scheduled booster dose, were immunocompromised due to underlying disease or medical treatment, or were pregnant. Demographic data and information on prior COVID-19 infection and vaccinations (Feb 1, 2020 onwards) were extracted from the medical records **(Table 1)**. The study was performed in line with the guidelines of the Declaration of Helsinki and the study protocol was approved by the Institutional Review Board of Qatar Biomedical Research Institute (ID 2022-52) and the Hamad Medical Corporation (ID MRC-

### Sample collection and processing

Peripheral blood was collected in 10ml EDTA blood tubes at three timepoints; at the time of the third dose (timepoint 1), 3-months after the third dose (timepoint 2) and 6-months post third dose (timepoint 3). Serum was collected after centrifugation at 3000 rpm for 10 minutes and stored at -80°C. Peripheral blood mononuclear cells (PBMCs) were isolated from freshly collected blood samples using SepMate™ density gradient centrifugation (85460; Stem Cell Technologies) according to the manufacturer’s guidelines. Next, isolated PBMCs were resuspended in freezing media (50% FBS, 40% serum-free Roswell Park Memorial Institute 1640 medium (RPMI), 10% Dimethyl sulfoxide) and stored in liquid nitrogen until further use.

### Enzyme-linked immunosorbent spot (ELISpot)

We used two distinct ELISpot assays to quantify the number of immune cells that secrete either IFN-γ (3420-4AST-P1-1; Mabtech, Nacka Strand, Sweden) or granzyme B (3486n-4APW-P1-1; Mabtech, Nacka Strand, Sweden) in response to a pool of SARS-CoV-2 peptides. Immune cell reactivity was measured against 166 peptides derived from the S1 domain of the spike protein (amino acids 13-685, divided into two peptide pools S1 and S2) and 47 synthetic peptides, covering the spike, nucleoprotein, membrane protein, ORF3a and ORF7a (SNMO peptide pool). ELISpot assays were conducted according to the manufacturer’s guidelines using 2.5x10e5 PBMCs/well in duplicate with a final concentration of 2 ug/ ml of each peptide. In addition, wells with PBMCs alone were used as negative control and PBMCs treated with an anti-human anti-CD3 antibody (mAb CD3-2, #3420-4HST-10, Mabtech, Nacka Strand, Sweden) overnight served as positive control. After 48 hours of incubation with the peptides at 37 °C, PBMCs were removed, the plates were washed, and spots were developed. To enable detection of secreted IFN-γ, plates were incubated with 7-B6-1-biotin detection antibody for 2 hours, followed by 1 hour incubation with Streptavidin-HRP and addition of TMB substrate.

Detection of granzyme B spot forming units (SFUs) was obtained using the MT8610-biotin detection antibody, Streptavidin-ALP and BCIP/NBT-Plus substrate according to the manufacturer’s instructions. In addition to the IFN-γ and granzyme B ELISpot assays, we also performed an IgG ELISpot assay to enumerate B cells that are secreting human IgG in response to the Sars-CoV-2 receptor binding domain (RBD) (3850-4HPW-R1-1; Mabtech, Nacka Strand, Sweden). Moreover, to gain insight into the magnitude and longevity of the humoral SARS-CoV-2 immune response of vaccinated individuals, PBMCs were pre-stimulated with R848 (1 μg/ml) and recombinant human IL-2 (10 ng/ml) for 3 days to promote the differentiation of memory B cells into antibody-secreting cells, enabling their quantification through measurement of IgG secretion. Next, pre-stimulated and unstimulated cells were seeded in duplicate at 2.5x10e5 cells/well in the ELISpot plate which was pre-coated with anti-human IgG monoclonal antibodies. After 48 hours at 37 °C, RBD-specific IgG spots were detected using a WASP-tagged RBD protein, followed by anti-WASP-HRP and TMB substrate. Finally, for each ELISpot assay the number of SFUs were determined using the AID iSpot ELISpot reader (Autoimmun Diagnostika GmbH, Strasburg, Germany). Representative images are depicted in **Fig S1-S3**.

### SARS-CoV-2 IgG/IgM enzyme-linked immunosorbent assay (ELISA)

In addition to the IgG ELISpot assay, secretion of SARS-CoV-2 specific IgG/IgM antibodies was determined using an in-house developed ELISA. In short, 96-well plates (Nunc, Maxisorp) were coated overnight at 4°C with 1μg/ml SARS-Cov-2 spike protein or Nucleoprotein in 0.2M NaHCO_3_ (pH 9.6). Plates were washed three times with PBST (0.05% Tween-20) and blocked for 1 hour at room temperature using PBST-2.25% gelatin. Next, diluted serum samples (1:800) were added to washed plates for 2 hours (room temperature, 100 rpm), followed by incubation with either goat anti-human IgG-HRP or goat anti-human IgM-HRP for 1 hour at room temperature. Finally, TMB substrate was added for 20min, and absorbance values were measured at 450nm using the EnVision® Multilabel Plate Reader (PerkinElmer).

### SARS-CoV-2 neutralizing antibody assay

To assess the presence of SARS-CoV-2 antibodies with neutralizing abilities, we utilized an in-house neutralization antibody (NAb) assay. Briefly, recombinant hACE2 protein (1μg/ml in 0.2 M NaHCO_3_, pH 9.6) was coated on 96-well ELISA plates (Maxisorp, Nunc) at 4°C overnight. Plates were washed three times with PBST (0.05% Tween-20) and blocked with 2.25% gelatin in PBST for 1 hour at room temperature. Serum samples were diluted (1:10) and preincubated with 100 ng/ml RBDmFc (Genscript) in blocking buffer for 1 hour at room temperature, after which they were added to the pre-coated ELISA plate for 1 hour. In parallel, anti-SARS nanobody NbS72-Biv (500 ng/ml) was preincubated with RBDmFc (Genscript) to serve as neutralization control. Next, wells were incubated with goat anti-mouse Fc-HRP (1:10,000) for 1 hour, followed by TMB substrate for 20min. Absorbance values were measured at 450nm using the EnVision® Multilabel Plate Reader (PerkinElmer).

### Immune cell phenotyping by flow cytometry

We characterized the presence of T and B cell subpopulations using DuraClone IM T cells (Beckman Coulter; #B53328) and DuraClone IM B cell tubes (#B53318, Beckman Coulter) respectively. DuraClone IM T cell tubes were supplemented with CD62L-SBV670 (#MCA1076SBV670, BioRad) and CD45RO-SBV790 antibodies (#MCA461SBV790, BioRad), while the B cell panel was expanded by adding CD56-PC5.5 (#A79388, Beckman Coulter) and CD16-APC-A700 (#B20023, Beckman Coulter) antibodies. A total of 3.0x10e5 PBMCs were resuspended in stain buffer (#554656, BD Pharmingen^TM^) and added to the DuraClone tubes, which were vortexed for 5 seconds and incubated for 15 min in the dark at room temperature. Next, the cells were washed in DPBS (#14190-144, Gibco) and resuspended in PBS prior to analysis on the LSRFortessaTM X-20 flow cytometer (BD Biosciences) using FACS Diva Software (BD Biosciences). For each sample, 30,000 events were recorded, and further analysis was performed using FlowJo™ Software (BD Biosciences, version 10.8).

### Statistical analysis

Statistical analyses were performed using GraphPad PRISM V9.5.1 (GraphPad Software, CA, USA). Data normality was assessed by Shapiro-Wilk test and differences between groups were analyzed using the unpaired t-test or one-way ANOVA test with Tukey correction. A p value ≤ 0.05 was considered significant.

## RESULTS

### Natural infection and booster vaccination differentially stimulate SARS-Cov-2 specific cellular immune responses

A total of 70 participants were enrolled in the study. For 45 study participants, we collected blood samples at three timepoints; prior to the third vaccine dose or booster (0M), three months post-booster (3M) and six months post-booster (6M). Among these participants, 18 had no prior PCR-confirmed SARS-CoV-2 infection before receiving the booster dose **(Fig 1 -top)**. These blood samples were used to assess cellular and humoral immune responses through ELISpot and flow cytometry analyses. An additional 25 participants, from whom less than three blood samples were obtained, were included in the study to evaluate anti-SARS-CoV-2 specific antibody levels, including the level of antibodies with neutralizing activity **(Fig 1 - bottom)**.

**Figure 1.**
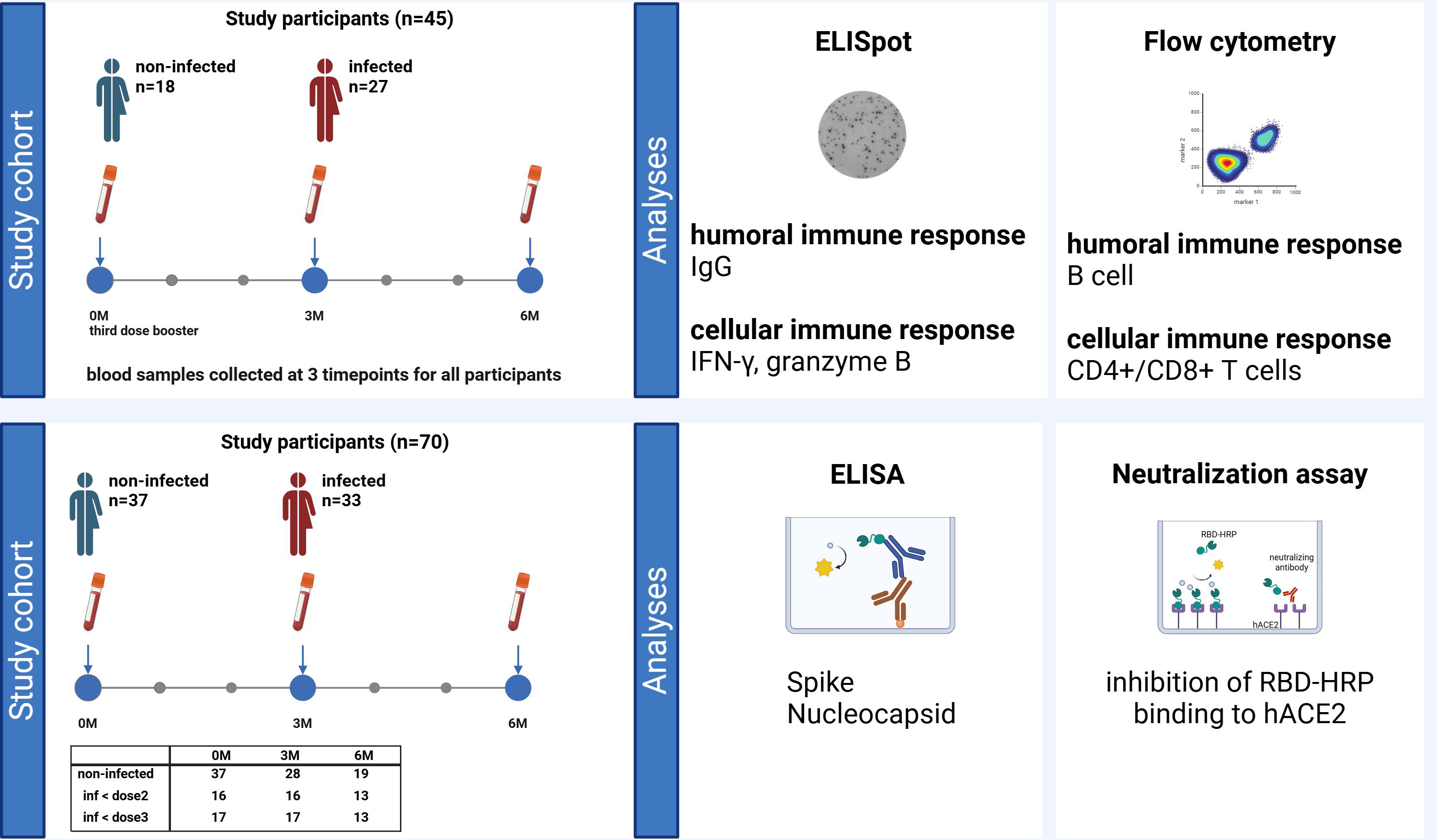
Study design diagram. Flowchart depicting the study cohort, blood sample collection and assays performed.

Comparative analysis of IFN-γ secretion following incubation of peripheral blood lymphocytes peptide pools) up to three to six months post-booster (3M, 6M) in participants who were previously infected with SARS-CoV-2. We found a significantly higher IFN-γ release in response to anti-spike peptides (S1 and S2 pool) in previously infected participants at 3-months and 6-months post-booster as compared to non-infected participants **(Fig 2A, Fig S1)**. This is likely the combined result of an elevated baseline, albeit non-significant, anti-spike response and the boosting effect of the third vaccine dose. In accordance, we observed a trend for increased anti-spike IFN-γ responses in previously infected participants following the administration of the third vaccine dose (S1 – 6M, S2 - 3M) **(Fig 2B)**. No differences in anti-SNMO IFN-γ responses were observed in relation to previous infection or booster vaccination **(Fig 2A-B)**. Next, we stratified the previously infected participants based on the timing of their infection relative to the second and third vaccine doses: “before dose 2”, or “before dose 3” for those infected between dose 2 and dose 3. Upon stratification, we observed higher anti-spike IFN-γ responses before administration of the third vaccine dose (0M - S1 and S2) in individuals with a recent SARS-CoV-2 infection (“before dose 3”) as compared to non-infected individuals or those infected “before dose 2” **(Fig 2C)**. Of note, booster vaccination did not increase anti-spike responses in our study cohort **(Fig 2D)**. In addition to elevated pre-booster anti-spike responses, recently “before dose 3” infected participants exhibited a strong baseline IFN-γ response against the SNMO peptide pool **(Fig 2C)**, however, following administration of the third vaccine dose this response seems to decrease over time to similar levels as observed in non-infected and “before dose 2” previously infected participants **(Fig 2D)**. Together, these findings suggest that individuals with a more recent natural SARS-CoV-2 infection exhibit stronger anti-spike and anti-SNMO T cell responses, which naturally taper off with time and are not further enhanced or sustained by booster vaccination.

**Figure 2.**
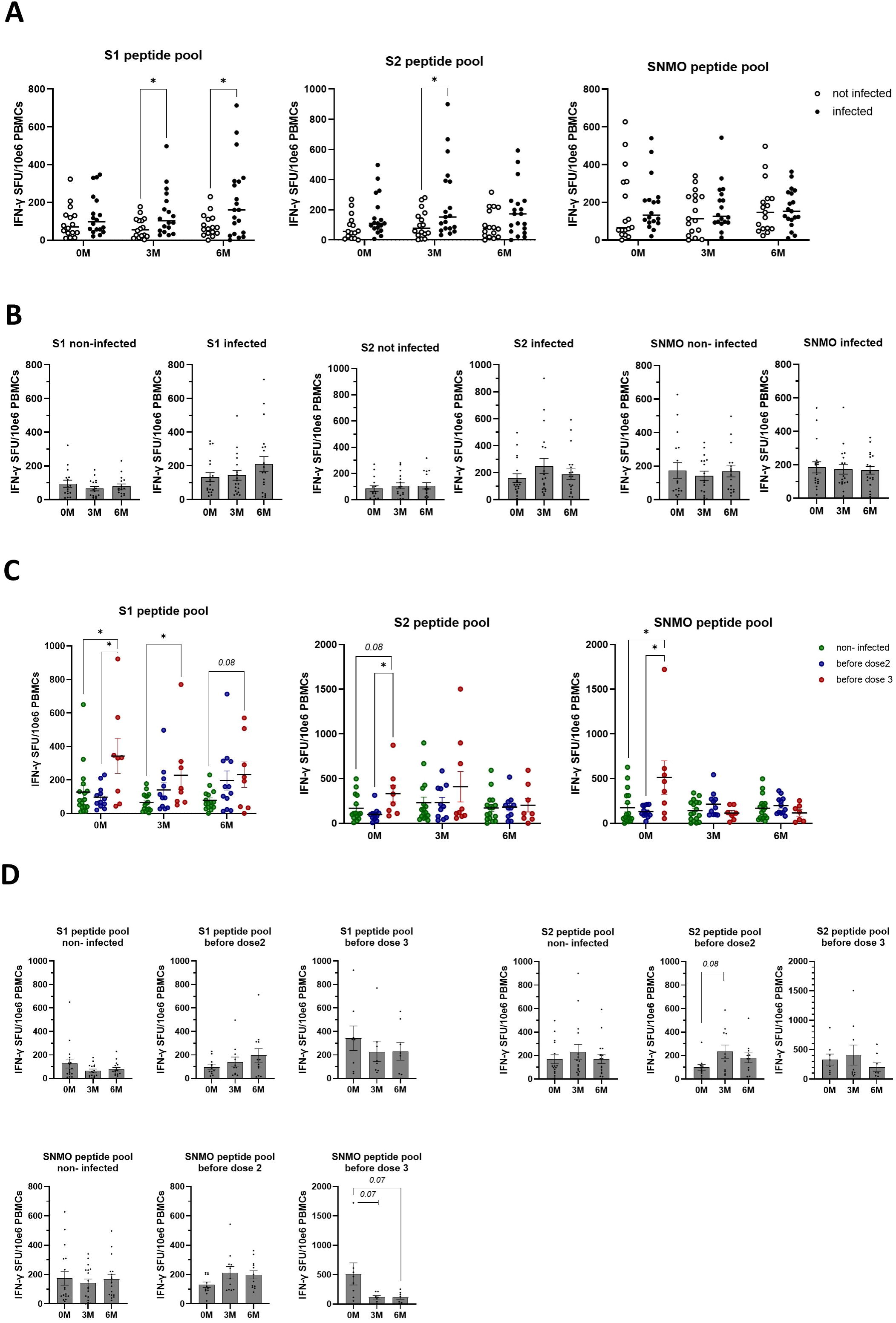
Natural exposure to SARS-CoV-2 enhances IFN-γ cellular immune responses. **A)** T cell responses against SARS-CoV-2 spike and SNMO peptide pools in non-infected and infected participants before and after booster vaccination, as determined by IFN-γ ELISpot analysis. **B)** Quantification of anti-spike and anti-SNMO specific IFN-γ responses within non-infected and infected participants pre- and post-booster vaccination. **C)** T cell responses against SARS-CoV-2 spike and SNMO peptide pools in non-infected and infected participants, stratified by the time of infection, as determined by IFN-γ ELISpot analysis of pre- and post-booster samples. **D)** Quantification of anti-spike and anti-SNMO specific IFN-γ responses within non-infected and stratified infected participants following booster vaccination. Scatter plots represent median, and bar charts depict mean with standard error of mean (±SEM). Statistical analysis performed using unpaired Student’s t-test or one-way ANOVA with Tukey correction. *p ≤ 0.05.

To evaluate the functional status of SARS-CoV-2 specific T cell responses, we assessed the effect of booster vaccination and pre-booster natural infection on the cytotoxic activity of CD8+ T lymphocytes by measurement of granzyme B release in response to exposure to the same SARS-CoV-2 peptide pools. At time of administration of the third vaccine dose, we did not observe any differences in the number of granzyme B-producing T cells in response to SARS-CoV-2 spike or SNMO peptides **(Fig 3A**, **Fig 3C, Fig S2)**. In contrast, an increase in anti-spike granzyme B responses, in particular anti-S2, was observed in non-infected participants following booster vaccination (S1 – 6M, S2 – 3M and 6M) **(Fig 3B)**. Furthermore, booster vaccination enhanced late anti-SNMO specific responses in previously infected participants (SNMO – 6M) **(Fig 3B)**. No differences were found when previously infected participants were stratified by the time of natural infection **(Fig 3C-D)**. Thus, natural SARS-CoV-2 infection appears to predominantly prime memory Th1 cell responses, characterized by IFN-γ release, whereas booster vaccination more likely drives cytotoxic cellular responses, marked by granzyme B production, particularly in the absence of prior natural infection.

**Figure 3.**
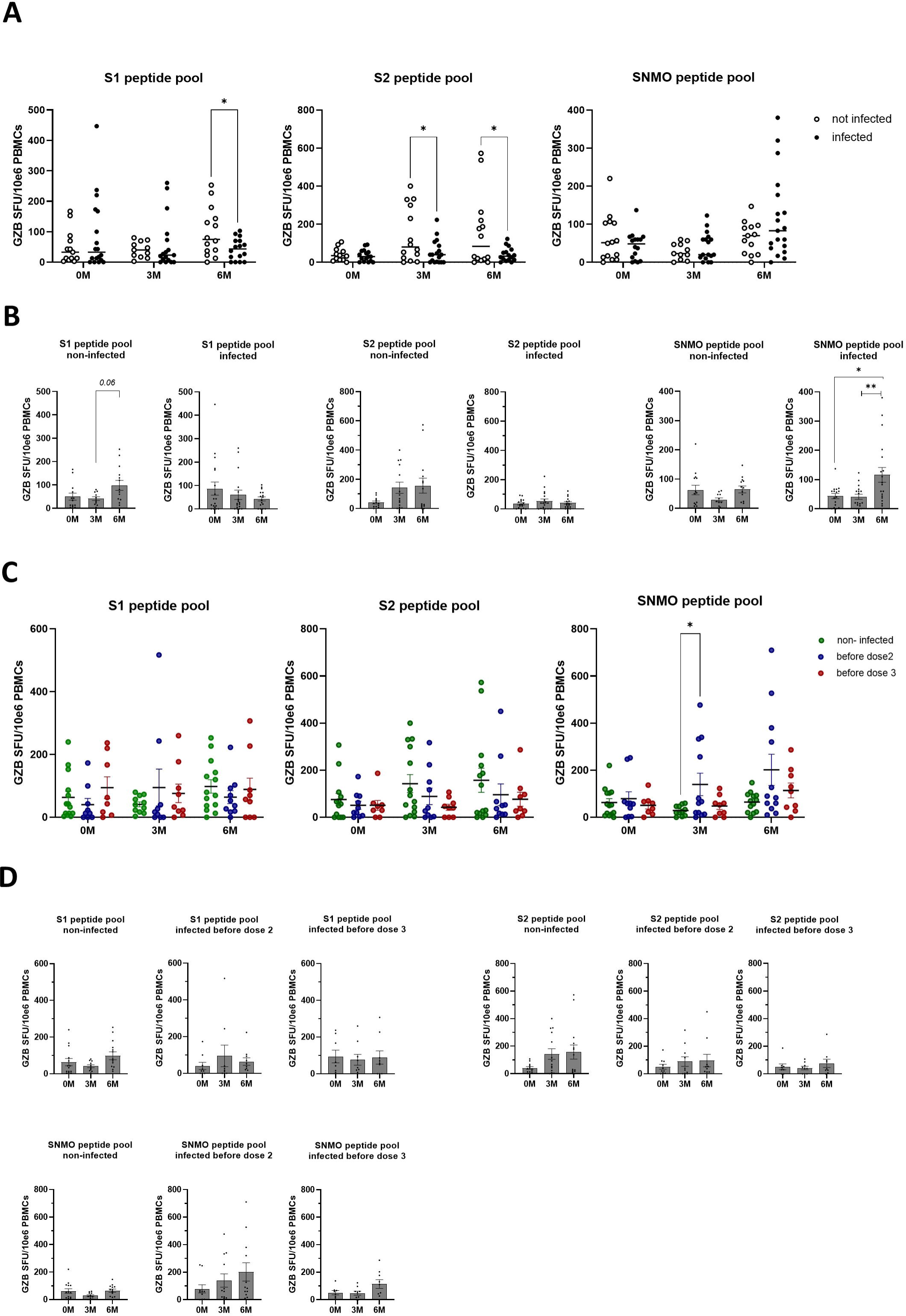
SARS-CoV-2 specific cytotoxic immune responses are enhanced by booster vaccination in the absence of prior natural infection. **A)** Cytotoxic T cell responses against SARS-CoV-2 spike and SNMO peptide pools in non-infected and infected participants before and after booster vaccination, as determined by granzyme B ELISpot analysis. **B)** Quantification of anti-spike and anti-SNMO specific granzyme B responses within non-infected and infected participants pre- and post-booster vaccination. **C)** Cytotoxic T cell responses against SARS-CoV-2 spike and SNMO peptide pools in non-infected and infected participants, stratified by the time of infection, as determined by granzyme B ELISpot analysis of pre- and post-booster samples. **D)** Pre- and post-booster quantification of anti-spike and anti-SNMO specific granzyme B responses within non-infected and previously infected participants, stratified by the time of infection. Scatter plots represent median, and bar charts depict mean with standard error of mean (±SEM). Statistical analysis performed using unpaired Student’s t-test or one-way ANOVA with Tukey correction. *p ≤ 0.05, ** p ≤ 0.01.

### Natural infection induces robust SARS-CoV-2 specific humoral responses with neutralizing abilities, which can be enhanced by booster vaccination

Based on our observations that SARS-CoV-2 natural infection and booster vaccination likely prime different components of the antiviral cellular immune response – specifically Th1 and cytotoxic responses - we set out to study their effects on SARS-CoV-2 specific humoral memory immune responses. Previously infected participants, particularly those with recent infections (“before dose 3”), displayed a higher number of IgG-secreting memory B cells before receiving the third vaccine dose **(Fig 4A-B, Fig S3)**. Furthermore, booster vaccination induced a transient increase in IgG-positive memory B cells with a peak at 3-months in non-infected participants followed by a decline by 6-months, while in “before dose 2” previously infected participants the number of IgG-positive memory B cells peaked by 6-months (**Fig 4A-B)**. This suggests that previously infected participants demonstrate a robust and durable humoral memory response which can be activated upon antigen re-exposure, and that booster vaccination can further enhance B cell responses. To validate these findings, we developed an ELISA to detect IgGs against the full-length spike protein. In accordance with our IgG ELISpot results, we observed a stronger baseline IgG-secreting memory B cell response in previously infected individuals, particularly in those with recent infections (“before dose 3”) **(Fig 4C-D)**. Furthermore, we confirmed that booster vaccination enhanced those responses in all participants **(Fig 4C-D)**. Further, we also observed a steady increase in anti-nucleocapsid IgG levels in recently infected participants following the third vaccine dose **(Fig 5A-B)**, further indicating the induction of a memory B cell response as a result of both natural infection and booster vaccination. To further assess the neutralizing abilities of these humoral responses, we used an in-house neutralization assay that demonstrated an increase in neutralizing antibody activity in infection-naïve and in previously infected participants post-booster, particularly in recently infected participants (“before does 3”), underscoring the effects of booster vaccination on memory B cell induction **(Fig 5C-D)**.

**Figure 4.**
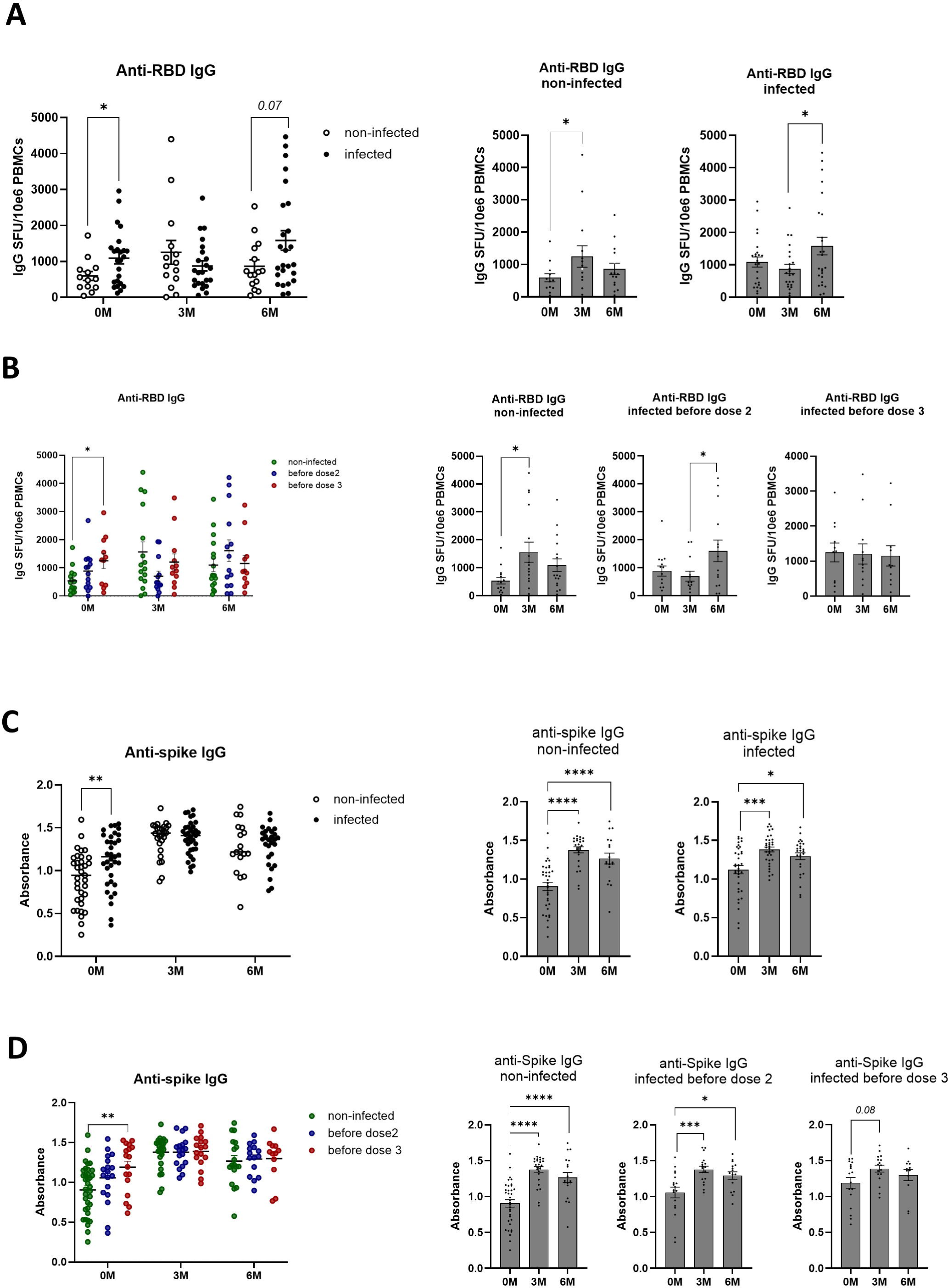
Effect of prior natural infection and booster vaccination on anti-SARS-CoV-2 RBD and spike IgG levels. **A)** Quantification of anti-RBD IgG producing memory B cells in non-infected and infected participants before and after booster vaccination, as determined by anti-RBD IgG ELISpot analysis. **B)** Quantification of anti-RBD IgG responses within non-infected and infected participants pre- and post-booster vaccination. **C)** Anti-spike IgG memory B cell responses in non-infected and infected participants, stratified by the time of infection, as determined by anti-spike IgG ELISpot analysis of pre- and post-booster samples. **D)** Quantification of anti-spike IgG memory B cell responses within non-infected and stratified infected participants following booster vaccination. Scatter plots represent median, and bar charts depict mean with standard error of mean (±SEM). Statistical analysis performed using unpaired Student’s t-test or one-way ANOVA with Tukey correction. *p ≤ 0.05, ** p ≤ 0.01, *** p ≤ 0.001, **** p ≤ 0.0001.

**Figure 5.**
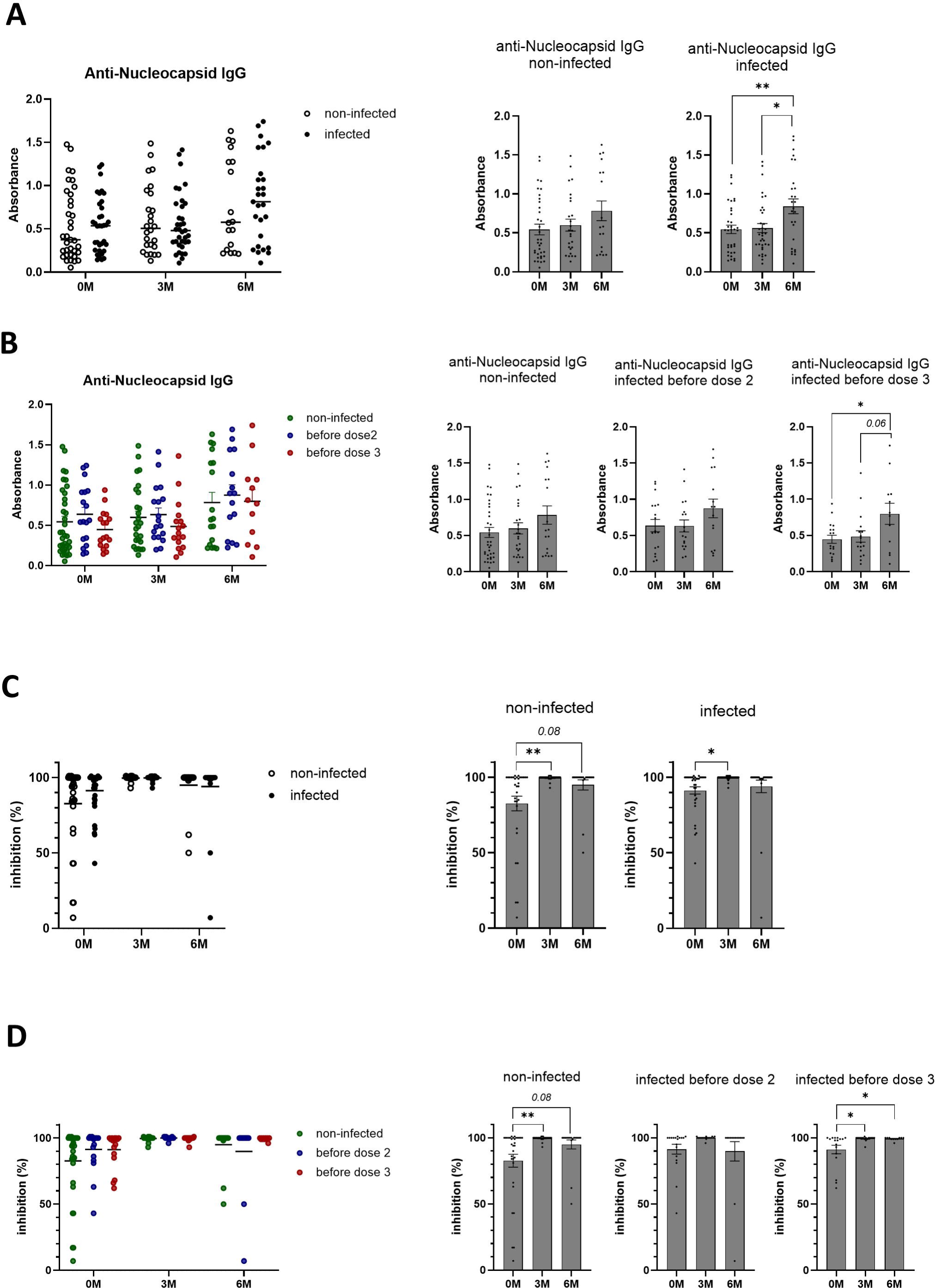
Booster vaccination increases the presence of SARS-CoV-2 neutralizing antibodies. **A)** Anti-nucleocapsid IgG levels in non-infected and previously infected participants pre- and post-booster vaccination, as determined by in-house ELISA. **B)** Anti-nucleocapsid IgG production in non-infected and infected participants, stratified by the time of infection, measured by in-house ELISA. **C)** Quantification of SARS-CoV-2 IgG with neutralizing ability in non-infected and previously infected participants pre- and post-booster vaccination, as determined by in-house neutralization assay. **D)** Analysis of SARS-CoV-2 neutralizing IgG antibodies in non-infected and infected participants, stratified by the time of infection, as measured by in-house ELISA. Scatter plots represent median, and bar charts depict mean with standard error of mean (±SEM). Statistical analysis performed using unpaired Student’s t-test or one-way ANOVA with Tukey correction. *p ≤ 0.05, ** p ≤ 0.01.

### Phenotyping analysis of cellular and humoral immune responses

Given the presence of a cytotoxic cellular and neutralizing humoral immune response in participants with previous natural infection and following booster vaccination, we further assessed the phenotype of the immune cells that provide this immunological memory **(Fig S4-S5)**. We did not observe any differences in CD8+ T cell phenotype that may complement the increased cytotoxic activity following natural infection and booster vaccination **(Fig 6A)**. In line with our observations demonstrating the presence of an enhanced Th1 immune response in previously infected participants, we found a decrease in the number of naïve CD4+ T cells (TN) following booster vaccination of recently infected participants who also exhibited a trend for a higher number of CD4+ effector memory cells (TEM) at baseline **(Fig 6B)**. Of note, we did not find any change in the number of CD4+ or CD8+ T cells expressing PD-1, suggesting that neither CD4+ nor CD8+ T cells exhibited an exhausted phenotype attributable to PD-1/PD-L1 signaling **(Fig 6C)**. Moreover, we observed a higher pre-booster number of class-switched B cells which was maintained for up to 6 months post-booster in recently infected individuals (“before dose 3”), underscoring the effect of natural infection on the development of a robust and durable memory B cell response **(Fig 6D)**.

**Figure 6.**
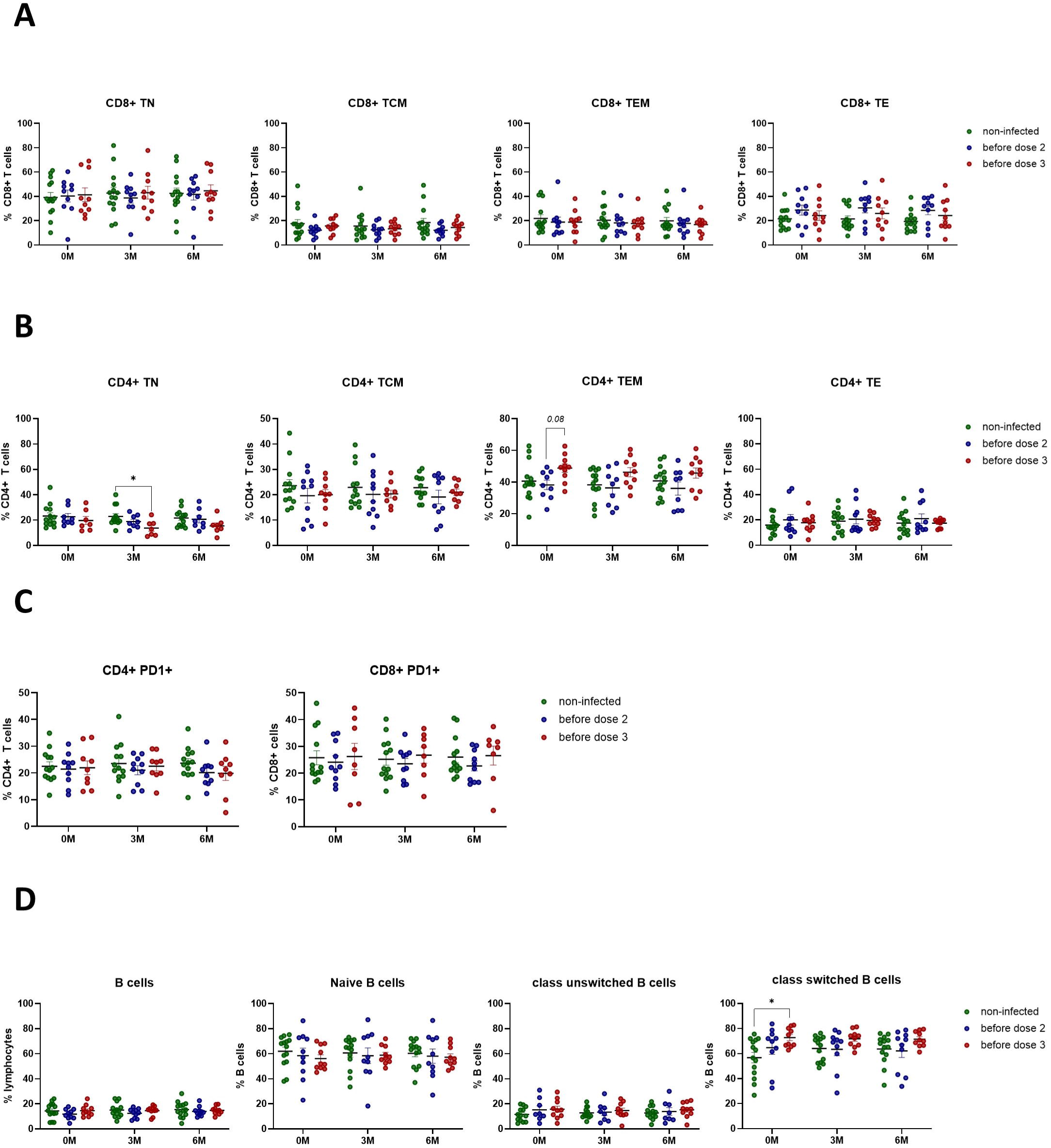
SARS-CoV-2 infection activates the anti-viral immune response by promoting CD4+ T effector memory and inducing class-switched B cell responses. **A)** Flow cytometry analysis of CD8+ T cell phenotypes. **B)** Flow cytometry analysis of CD4+ T cell phenotypes. **C)** Flow cytometry analysis of PD1 expression on CD4+ and CD8+ T cells. **D)** Flow cytometry analysis of B cell phenotypes. Scatter plots represent mean with standard error of mean (±SEM). Statistical analysis performed using unpaired Student’s t-test or one-way ANOVA with Tukey correction. *p ≤ 0.05. TN, naïve; TCM, central memory; TEM, effector memory; TE, effector T cells.

## DISCUSSION

In March 2020, the WHO declared COVID-19 a pandemic, making it the first pandemic caused by a coronavirus. Given its high infection and mortality rate, worldwide efforts focused on limiting the spread of the SARS-CoV-2 virus through implementation of nation-wide vaccination campaigns. Following the initial success of single- and two-dose vaccine approaches, vaccine effectiveness waned over time, raising the question whether a third vaccine dose or booster could mitigate the decline in protection. Here, we present a comprehensive, longitudinal analysis of cellular and humoral immune responses against SARS-CoV-2 pre- and post-booster vaccination and provide insights into the role of prior natural infection in establishing immunological memory **(Fig 7)**.

**Figure 7.**
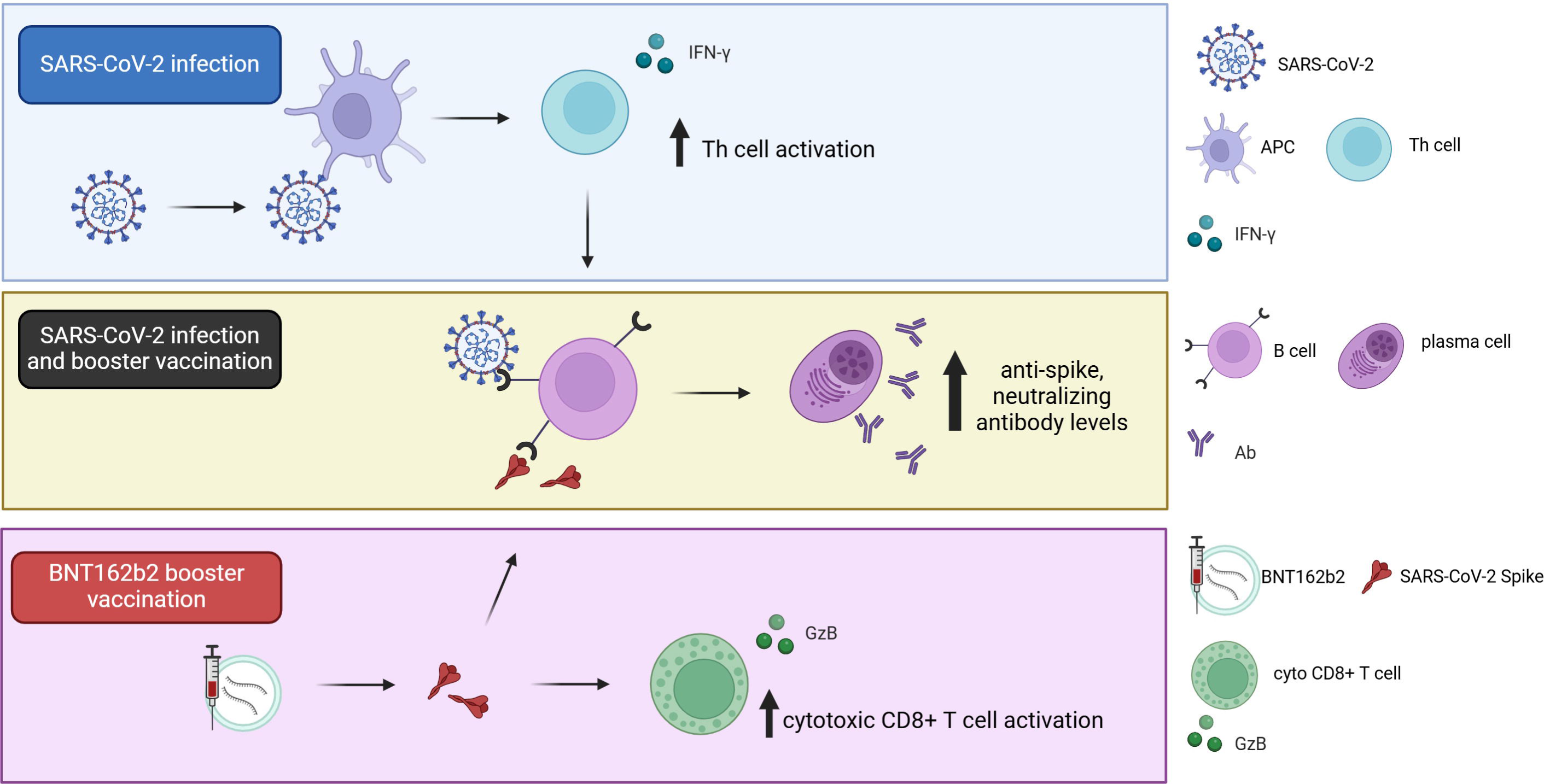
Diagram depicting the effect of SARS-CoV-2 natural infection and BNT162b2 third dose vaccination on cellular and humoral immune responses. Prior natural infection induces Th1 cellular and humoral immune responses. Booster vaccination with BNT162b2 further enhances humoral immune responses, and promotes a cytotoxic CD8+ T cell response in infection-naïve individuals.

In our study, we demonstrated that previously infected participants exhibited robust humoral immune responses, characterized by higher pre-booster anti-spike antibody levels, including those of neutralizing antibodies, which were further enhanced by booster vaccination. In addition, we found an increased number of class-switched B cells in the most recently infected participants prior to receiving the third vaccine dose. Non-infected or infection-naïve participants exhibited a sustained increase in the number of anti-spike, neutralizing antibodies following booster vaccination. Notably, this increase reached levels similar to those observed in previously infected individuals. These findings are in line with a Danish study that reported prolonged anti-spike antibody persistence following booster vaccination with BNT162b2 or mRNA-1273 (24). Furthermore, although previously infected participants showed a stronger baseline anti-spike humoral response than non-infected participants, similar responses were found following the booster vaccination, corroborating the previous study by Andrejko et al that reported similar vaccine effectiveness at 7-months post-vaccination independent of pre-vaccination naturally acquired immunity (19).

To further explore the anti-viral immune response, we also assessed any changes in the cellular immune response. Of note, we found that different arms of the cellular immune response were stimulated by either naturally acquired or vaccination-induced immunity. Recent exposure to a natural SARS-CoV-2 infection primarily induced a durable Th1 immune cellular response with increased IFN-γ production and a trend towards an increased number of CD4+ T effector memory cells. On the other hand, booster vaccination more readily induced the cytotoxic cellular immune response, marked by enhanced granzyme B release, in non-infected participants for up to 6-months after receiving the third vaccine dose. We did not find any increase in the number of PD-1 positive CD4+ or CD8+ T cells in our study participants, reflecting a lack of T cell exhaustion up to 6-months following the third vaccine dose. This raises the question whether cellular immune responses could be maintained for more than 6-months post-booster. Moreover, it would be of interest to study additional immune checkpoint markers as well as the exhaustion status of distinct CD4+ and CD8+ T cell subpopulations such as the T effector memory cells in future studies. Collectively, our findings indicate that prior natural SARS-CoV-2 infection and booster vaccination both stimulate the humoral immune response while activating different aspects of the cellular immune response, and that these responses can persist for at least 6-months after receiving the third vaccine dose of BNT162b2.

In conclusion, we demonstrated that booster vaccination of both infection-naïve and previously infected individuals plays a critical role in enhancing the anti-viral humoral and cellular immune responses and hence likely provides a broader protection in the population. Additional studies are warranted to explore the magnitude, longevity and exhaustion status of the immunological memory beyond 6-months post-booster.

## ACKNOWLEDGEMENTS

This work was supported by a grant from the Qatar Biomedical Research Institute (QB10-IDRP-2021), Qatar Foundation awarded to Dr Julie Decock.

## AUTHOR CONTRIBUTIONS

Remy Thomas processed samples, performed experiments, and wrote the first draft. Ahmed Zaqout recruited participants, collected samples and clinical data. Bakhita Meqbel processed samples performed some experiments and wrote the first draft. Umar Jafar performed some experiments and analyzed data. Nishant Vaikath performed some experiments. Abdullah Aldushain collected clinical data. Adviti Naik, Hibah Shaath, Neyla S Al-Akl, Abdi Adam, Houda Y Ali Moussa, Kyung C Shin and Rozaida Z Taha processed samples. Mohammed Abdukhattab and Muna A Almaslamani contributed to recruitment of participants. Nehad M Alajez, Abdelilah Arredouani, Yongsoo park, Sara A Abdulla, and Omar MA El-Agnaf revised the manuscript. Ali S Omrani conceptualized the study design and revised the manuscript. Julie Decock conceptualized the study, was responsible for funding acquisition and project administration, analyzed and visualized data, and wrote the draft manuscript. All authors read and approved the final manuscript.

## SUPPLEMENTARY MATERIAL

**Figure S1. IFN-γ release following administration of BNT162b2 third vaccine dose in non-infected and previously infected participants.** Representative images of IFN-γ ELISpot assays against SARS-CoV-2 S1, S2 and SNMO peptide pool. Activated peripheral blood lymphocytes, using anti-CD3 antibodies, were used as positive controls and cells without activation nor exposure to peptide pools served as negative controls. 0M, 0 months after third vaccine dose/baseline; 3M and 6M, 3 and 6 months after third vaccine dose.

**Figure S2. Granzyme B release following administration of BNT162b2 third vaccine dose in non-infected and previously infected participants.** Representative images of GzB ELISpot assays against SARS-CoV-2 S1, S2 and SNMO peptide pool. Activated peripheral blood lymphocytes, using anti-CD3 antibodies, were used as positive controls and cells without activation nor exposure to peptide pools served as negative controls. 0M, 0 months after third vaccine dose/baseline; 3M and 6M, 3 and 6 months after third vaccine dose.

**Figure S3. Anti-RBD IgG antibody levels following administration of BNT162b2 third vaccine dose in non-infected and previously infected participants.** Representative images of IgG ELISpot assays against SARS-CoV-2 RBD. Peripheral blood lymphocytes were stimulated with R848 and recombinant human IL-2 for 3 days prior to the assays to promote the differentiation of memory B cells into antibody-secreting cells. 0M, 0 months after third vaccine dose/baseline; 3M and 6M, 3 and 6 months after third vaccine dose.

**Figure S4. CD4+ and CD8+ immunoprofiling gating strategy.** CD4+ and CD8+ T cells were gated from CD3+ T cells, following gating for single cells and leukocytes, and analyzed for PD1+ expression or divided into four subpopulations using CD45RA and CCR7; naïve T cells (TN), central memory T cells (TCM), effector memory T cells (TEM) and effector T cells (TE).

**Figure S5. B cell immunoprofiling gating strategy.** CD19+ B cells were gated from single cell leukocytes and identified as naïve B cells using CD27 and IgD. Class unswitched and class switched memory B cells were identified using subgating of the IgM-IgD- and IgM+IgD+ populations based on CD27 and CD38 expression.

