## Supplementary figures and images for "Longitudinal cellular and humoral immune responses following Covid-19 BNT162b2-mRNA-based booster vaccination of craft and manual workers in Qatar"

### Fig S1

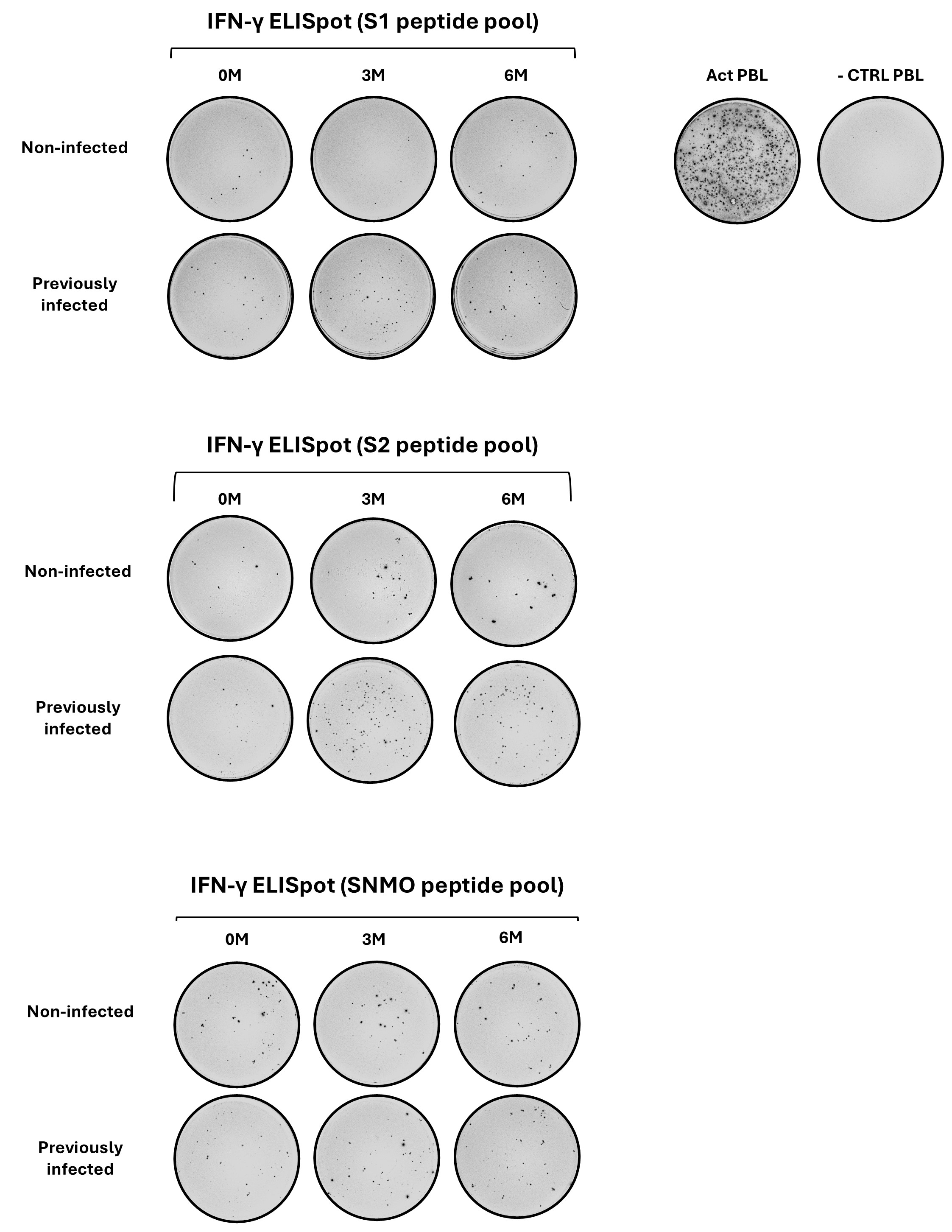

### Fig S2

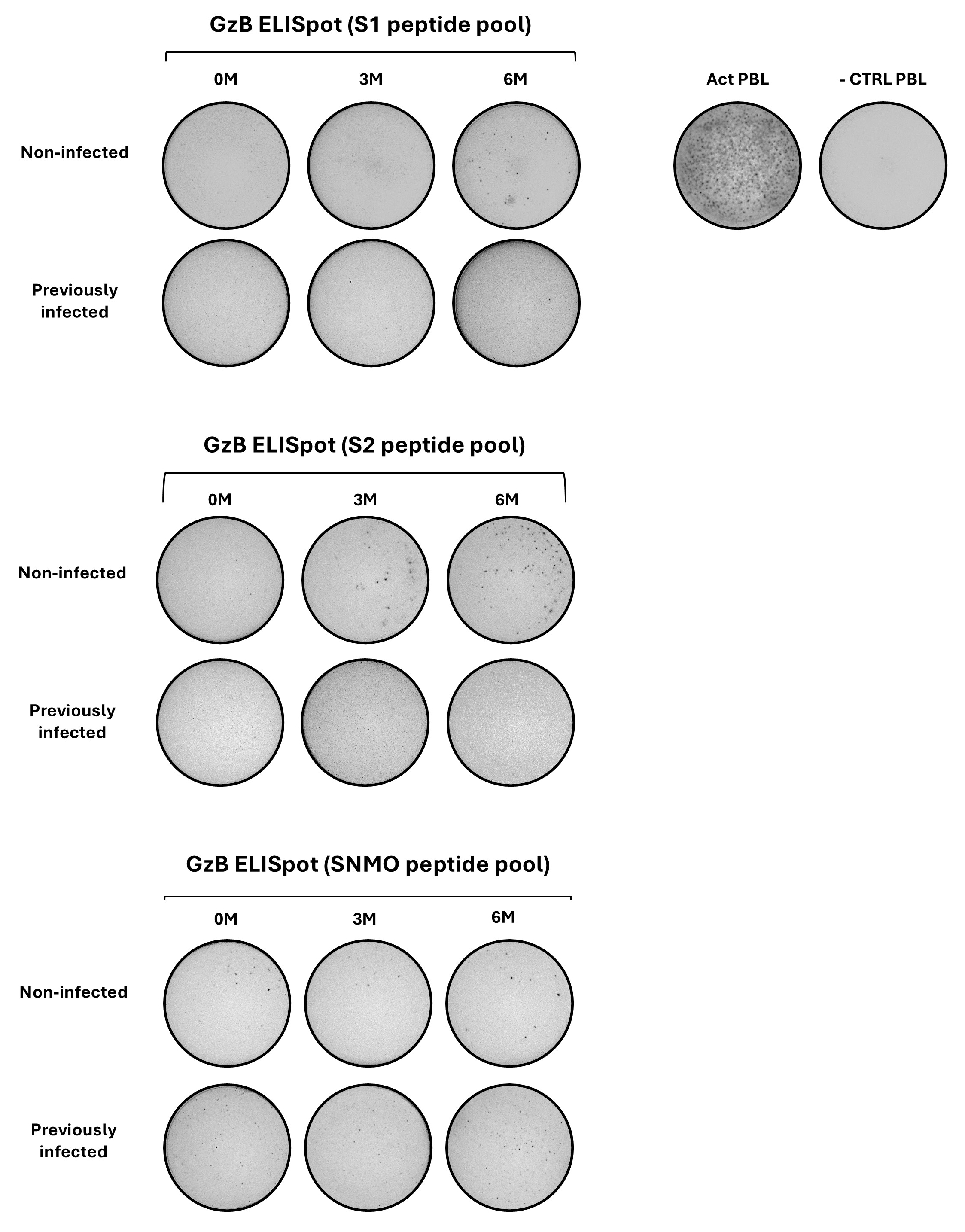

### Fig S3

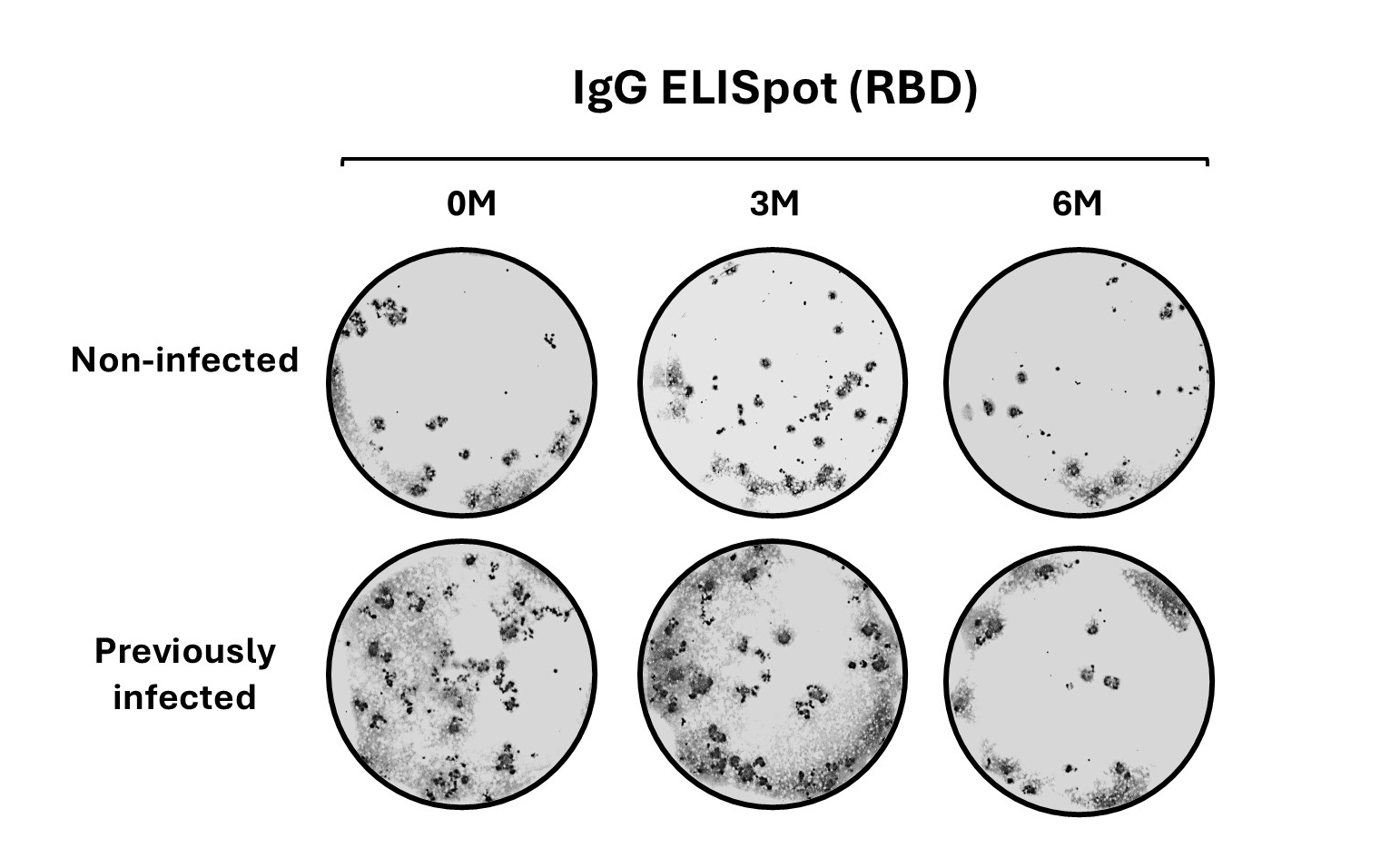

### Fig S4

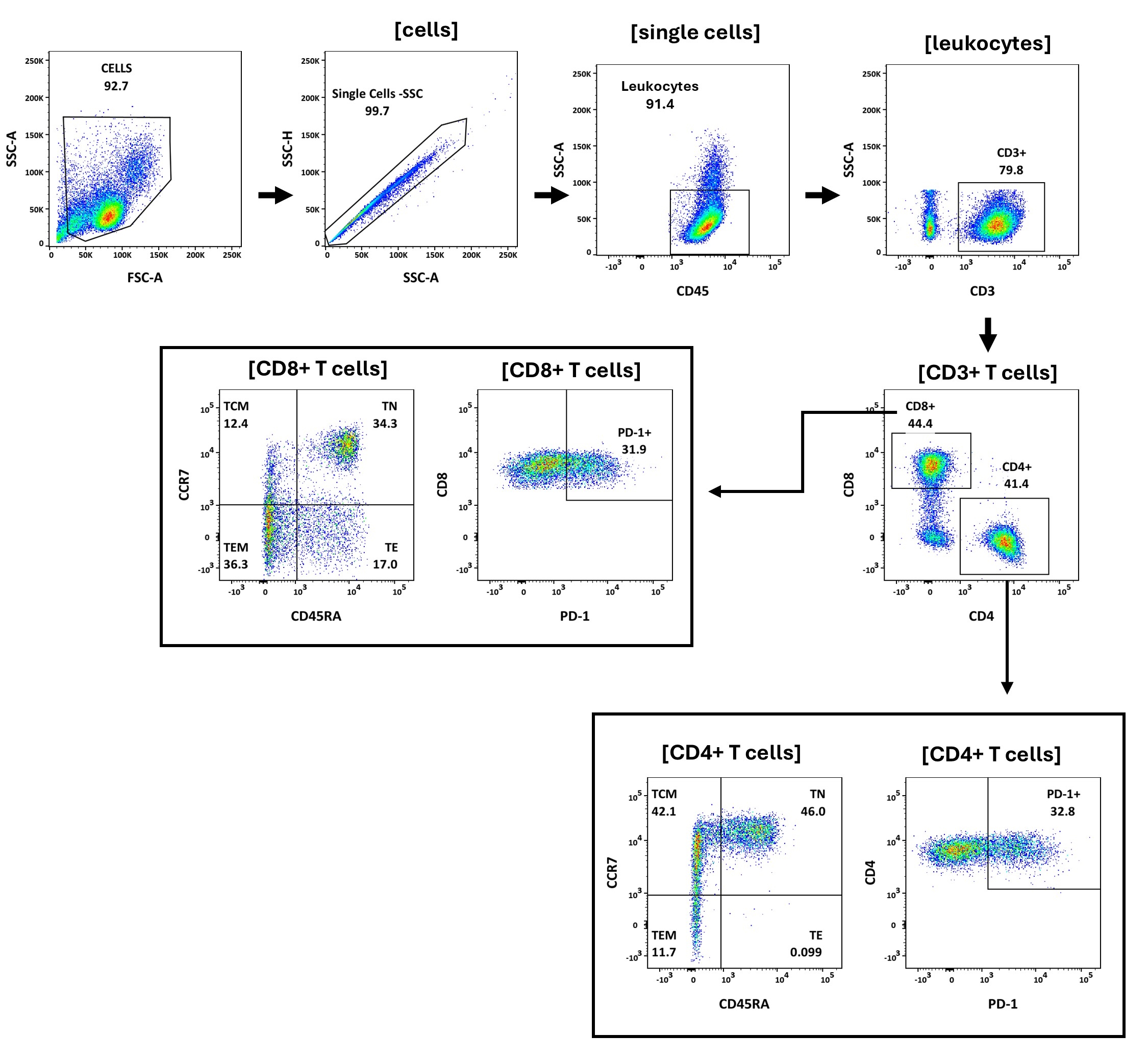

### Fig S5

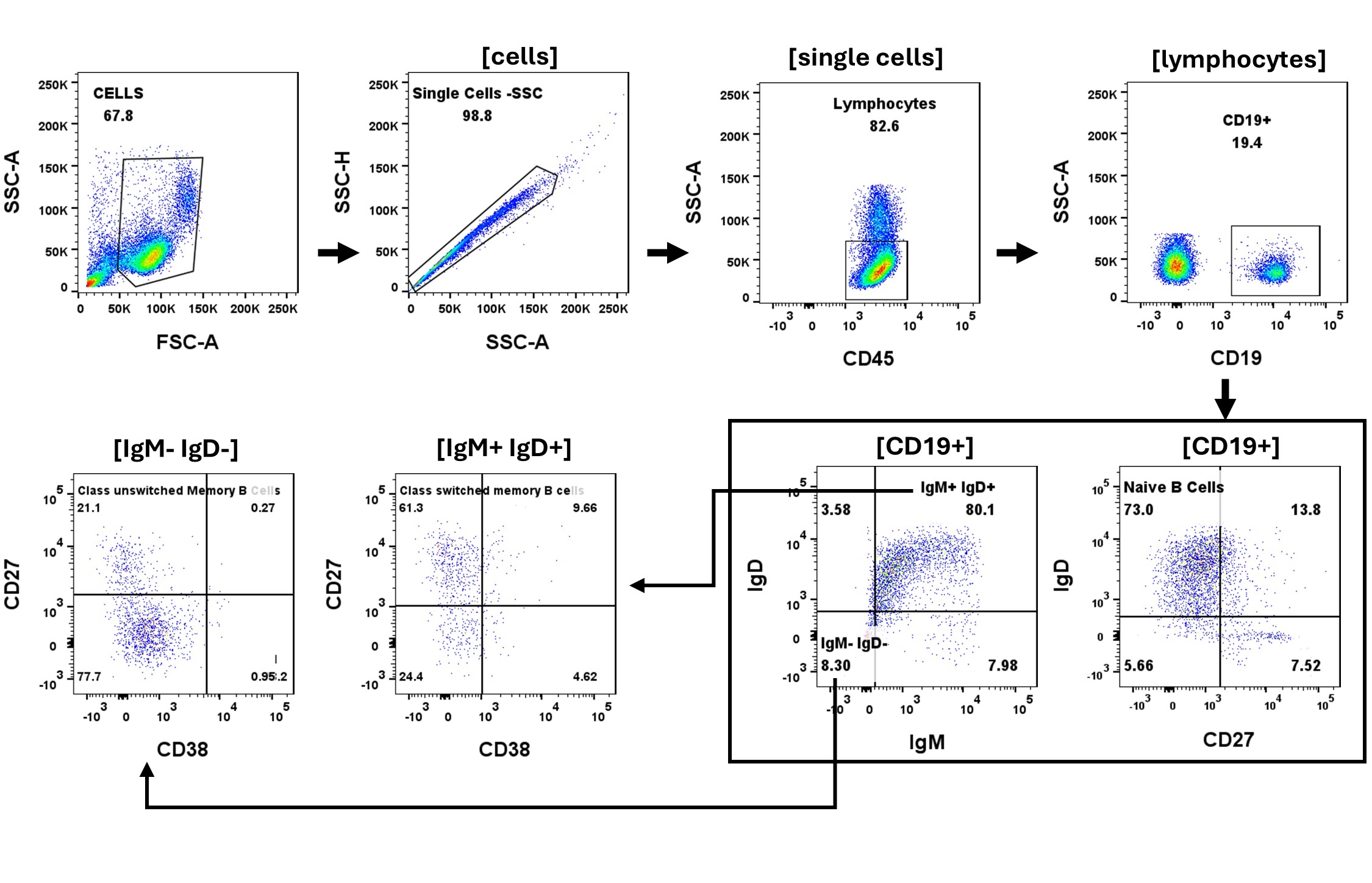
